# Vulture: Cloud-enabled scalable mining of microbial reads in public scRNA-seq data

**DOI:** 10.1101/2023.02.13.528411

**Authors:** Junyi Chen, Danqing Yin, Harris Y.H. Wong, Xin Duan, Ken H.O. Yu, Joshua W. K. Ho

## Abstract

The rapidly growing collection of public single-cell sequencing data have become a valuable resource for molecular, cellular and microbial discovery. Previous studies mostly overlooked detecting pathogens in human single-cell sequencing data. Moreover, existing bioinformatics tools lack the scalability to deal with big public data. We introduce Vulture, a scalable cloud-based pipeline that performs microbial calling for single-cell RNA sequencing (scRNA-seq) data, enabling meta-analysis of host-microbial studies from the public domain. In our scalability benchmarking experiments, Vulture can outperform the state-of-the-art cloud-based pipeline Cumulus with a 40% and 80% reduction of runtime and cost, respectively. Furthermore, Vulture is 2-10 times faster than PathogenTrack and Venus, while generating comparable results. We applied Vulture to two COVID-19, three hepatocellular carcinoma (HCC), and two gastric cancer human patient cohorts with public sequencing reads data from scRNA-seq experiments and discovered cell-type specific enrichment of SARS-CoV2, hepatitis B virus (HBV), and *H. pylori* positive cells, respectively. In the HCC analysis, all cohorts showed hepatocyte-only enrichment of HBV, with cell subtype-associated HBV enrichment based on inferred copy number variations. In summary, Vulture presents a scalable and economical framework to mine unknown host-microbial interactions from large-scale public scRNA-seq data. Vulture is available via an open-source license at https://github.com/holab-hku/Vulture.

## Introduction

Pathogenic diseases are considered a significant threat to global health, such as severe acute respiratory syndrome coronavirus 2 (SARS-CoV-2) in coronavirus disease 2019 (COVID-19), hepatitis B virus (HBV) and hepatitis C virus (HCV) in hepatocellular carcinoma (HCC) [1], and *H. pylori* in gastric cancer (GC) [2]. Single-cell or single-nucleus RNA sequencing (sc/snRNA-seq) has reformed the investigation of complex diseases and contributed to discoveries of host-microbial interaction mechanisms [3]–[7]. Due to the rapidly maturing scRNA-seq technologies, the exponentially growing public scRNA-seq data resources have become a gold mine for conducting *in silico* investigations toward host-microbial interactions.

In the current practice of scRNA-seq data processing, a key concern is the selection of the reference genomes when quantifying the reads. Most studies only align reads to the host genome or focus on limited microbial genomes [8]–[12]. This practice systematically risks missing either the known or unknown host-microbial interactions in the datasets. It is therefore worthwhile to perform re-analyses of existing public scRNA-seq data on the cloud to uncover the breadth of these interactions. According to the Human Cell Atlas [13] Data Portal, as of Dec. 2022 there are an estimated 12.3M cells from 2,400 specimens, totaling 38.1 TB in filesize of published human cellular droplet-based scRNA-seq data. As more and more files become available, cloud computing becomes increasingly enticing as the choice for performing large-scale re-analyses that can leverage huge amounts of computational resources without the need to purchase or maintain expensive hardware and avoid the transfer of large amounts of data.

Several tools have been developed for the identification of microbial reads in human scRNA-seq data on local machines. Viral-Track [14] is an existing computational pipeline that detects viral-host interactions in droplet scRNA-seq data by scanning host-unmapped reads for the presence of viral RNA. Based on a similar schema, Zhang et al. and Lee et al. developed PathogenTrack [15] and Venus [16] which have added capabilities. PathogenTrack can quantify bacteria in addition to viruses; the tool was benchmarked to be mostly correlated in microbial unique molecular identifiers (UMIs) called as and faster in run time than Viral-Track [15].Venus identifies viruses only but has another module to discover viral integration sites. However, due to the number of steps and certain tool choices in these pipelines, their scalability can still be improved.

These off-the-shell microbial calling methods for scRNA-seq are developed as command-line tools running on local computing environments, which may eventually struggle with the scale of published data on the cloud. Only with cloud computing can we obtain the scRNA-seq big data as well as leverage a huge amount of computational resources without the maintenance of expensive devices. Previously, we [17] developed the cloud-based Falco framework for scalable scRNA-seq analysis. On two public scRNA-seq datasets, it was 2.6-145.4 times faster than running on the local computing environments. Li et al. [18] developed a scalable scRNA-seq analysis framework based on the Terra platform called Cumulus afterward. With Cumulus, Delorey et al. [19] performed COVID-19 scRNA-seq dataset analysis on 420 specimens from 11 organs in 2021. In 2022, Edgar et al. [20] developed the Serratus framework. They reviewed 5.7 million transcriptome sequencing (RNA-seq) data for RNA-dependent RNA polymerases and identified more than 105 novel RNA viruses. However, Falco has supported an insufficient amount of up-to-date scRNA-seq protocols since 2017. Also, neither the Cumulus nor the Serratus pipeline focus on the host-microbial scRNA-seq analysis.

To perform a large-scale meta-analysis of public scRNA-seq data, we developed Vulture, which to our knowledge is the first cloud-based scalable framework for discovering microbial reads in public scRNA-seq data. It can be executed either on the cloud container services in parallel or a local environment. Our tool provides an easily modifiable, host-microbial combined reference that standardizes the gene transcript annotations of human and known human-host viruses and bacteria. Additional features of our Vulture are the support of multiple formats of raw sequencing file inputs, and the quality control metrics of the identified intracellular microbial UMIs. We benchmarked the scalability and cost-effectiveness of our tool, and show it outperforms existing solutions. With Vulture, we re-analyzed cohorts of COVID-19, hepatocellular carcinoma, and gastric cancer with public raw sequencing data of droplet scRNA-seq, and examined the host-microbial interactions of SARS-CoV-2, HBV, and *H. pylori*, respectively. Specifically, we detected an upregulation of chemokine receptor crosstalk along with the co-infection of SARS-CoV2 and human metapneumovirus (hMPV) from a COVID-19 bronchoalveolar lavage fluid (BALF) sample and a potential HBV-induced copy number variations from an HCC sample. The result shows the utility of viral calling to the full set of known host microbes.

## Materials and Methods

### Cloud infrastructure of Vulture on AWS batch infrastructure with Nextflow

The cloud framework of Vulture is described in **Supplementary Fig. S1a**. Vulture applications on the cloud are constructed in Docker containers. A container is a lightweight software unit that packages all our procedures and dependencies for sequence alignment, quality control, and downstream analysis. Containerized Vulture applications are managed by the Amazon Web Services (AWS) Batch service. AWS Batch is a batch management capability to efficiently run a huge amount of batch computing jobs on AWS. The Batch is a job scheduler composed of four elements including Compute Environments, Job queues, Job definitions, and Jobs. The Compute Environment specifies the computational resources required for a type of task. We applied the SPOT_CAPACITY_OPTIMIZED allocation strategy in Batch to prioritize the use of spot instances in Compute Environment. The Job Queue maps the Vulture pipeline task to one or more Compute Environments. The Job Definition is a template that assigns the Docker image to be employed in running a particular task along with its parameters such as the number of CPUs, the amount of memory, and other configurations. The Jobs binds a Job Definition to a specific Job Queue and executes the task command in the Docker container. In the Vulture pipeline, Job definitions and execution of Jobs are controlled by Nextflow, a language that streamlines the deployment of workflows on the commercial cloud and clusters. Nextflow creates the required Job Definitions and Jobs as needed. Each Job can use a different queue and Docker image. The Vulture container is published in DockerHub and Elastic Container Registry (ECR) that are accessible from the instances run by Batch. The Simple Storage Service (S3) bucket is where the input, output, and working directory of the Vulture pipeline are stored during execution.

### Construction of host-microbe combined reference genome

The first step of Vulture is to construct reference genomes and corresponding annotations for the host (human, in this study) and host-infection viruses and bacteria. We use 245 distinct human-host prokaryotes curated by the NCBI Genome [21] (https://www.ncbi.nlm.nih.gov/genome/browse#!/prokaryotes/) and 529 viral species from viruSITE [22] which together with the human reference genome hg38 form a combined reference set. The set is a collation of the reference genome fasta sequences and exon/transcript/gene gtf annotations of all species used. Non-host exons with a minimap2 [23] alignment to the host genome were removed due to ambiguity. The combined host-microbe reference genome was indexed using the ***genomeGenerate*** module from the STAR [24] tool.

### Quantifying reads from scRNA-seq data to count matrices

Vulture supports a variety of alignment algorithms to quantify scRNA-seq sequences with the constructed combined reference genome. Users can select STARsolo [24] (default), Cell Ranger [25], kallisto | bustools [26], and Alevin [27]. Sequence data can be quantified as a two-dimensional Unique Molecular Identifiers (UMI) count matrix of cells × genes and the corresponding Binary Sequence Alignment Map (BAM). BAM files are only generated if STARsolo or Cell Ranger is selected.

### Quality control of the mapped microbial reads and count matrix

After obtaining the results of the sequence alignment, we also perform additional quality control steps to increase the likelihood that the viral sequences found are intracellular and reliable. Vulture utilizes the EmptyDrops [28] algorithm to filter out the droplets with non-cellular ambient RNA. We then optionally perform various quality analyses on the BAM files documenting the sequence alignments, including multi-mapping of the resulting sequences to host or non-host genes and reads dispersion, which is the extent of unique positions the reads are aligned on a given transcript.

### Downstream analysis of scRNA-seq samples

The meta-analysis of the Vulture processed results is composed of several procedures. We applied SCANPY [29] for the COVID-19 sample and Seurat [30] for HCC and GC samples to perform scRNA-seq processing, and clustering, respectively. For the batch effects removal across different cohorts, we applied BBKNN [31] for the COVID-19 sample and Harmony [32] for HCC and GC samples. CellChat [33] is applied to calculate the ligand-receptor interactions among the annotated cell types. Copy number variation (CNV) inference and clone identification for the HCC sample is analyzed by the inferCNV [34] package.

### Cell-type enrichment of microbial UMI

We follow the idea in [19] to calculate the cell-type-specific enrichment score of intracellular microbes. The reason is that the number of microbial UMIs is small and their differences across cell types are difficult to observe. The enrichment score for cluster C in the clustering of cells is computed as follows:

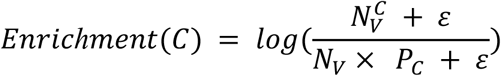

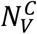 is the number of microbe positive cells in cluster C, 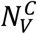 is the number of microbe positive cells in the whole cohort, *P_c_* is the proportion of the total number of cells in cluster C out of the total number of cells in the cohort, and *ε* is a small float to avoid zero subtractions.

The p-value of cell-type-specific enrichments of intracellular microbes was calculated by randomly permuting the identical number of microbe’s positive annotations to all cell types 10,000 times. The empirical p-value is the proportion of permutations that get an enrichment score not less than the actual score in the cohort out of 10,000 times. We also perform the FDR correction on the empirical p-value.

## Results

### Vulture: a cloud-based microbial calling framework for public scRNA-seq data

The architecture of Vulture is shown in **Fig. 1**. Vulture is composed of a bioinformatics analysis container, a cloud platform, and a workflow management tool. The container defines five main processes for performing microbe calling for sc/snRNA-seq data: sequence data retrieval, human-microbe combined reference construction (optional), reads alignment, quality control, and downstream analysis (optional). Detailed implementations are listed in the Method section and **Supplementary Fig. S1a**. Vulture receives two major inputs by default: 1) the sequencing files, and 2) microbe genome files. The input sequencing files can be a set of run accession numbers (prefixed by SRR) from SRA, a set of Amazon S3, or HTTP download URLs. Both fastq and bam files are supported. As for the input of combined reference, we provide a default host-microbe reference covering human and all human-host microbe genomes. Users can also build their custom combined genome by inputting a list of microbe genome accession numbers based on viruSITE or NCBI.

**Fig. 1.**
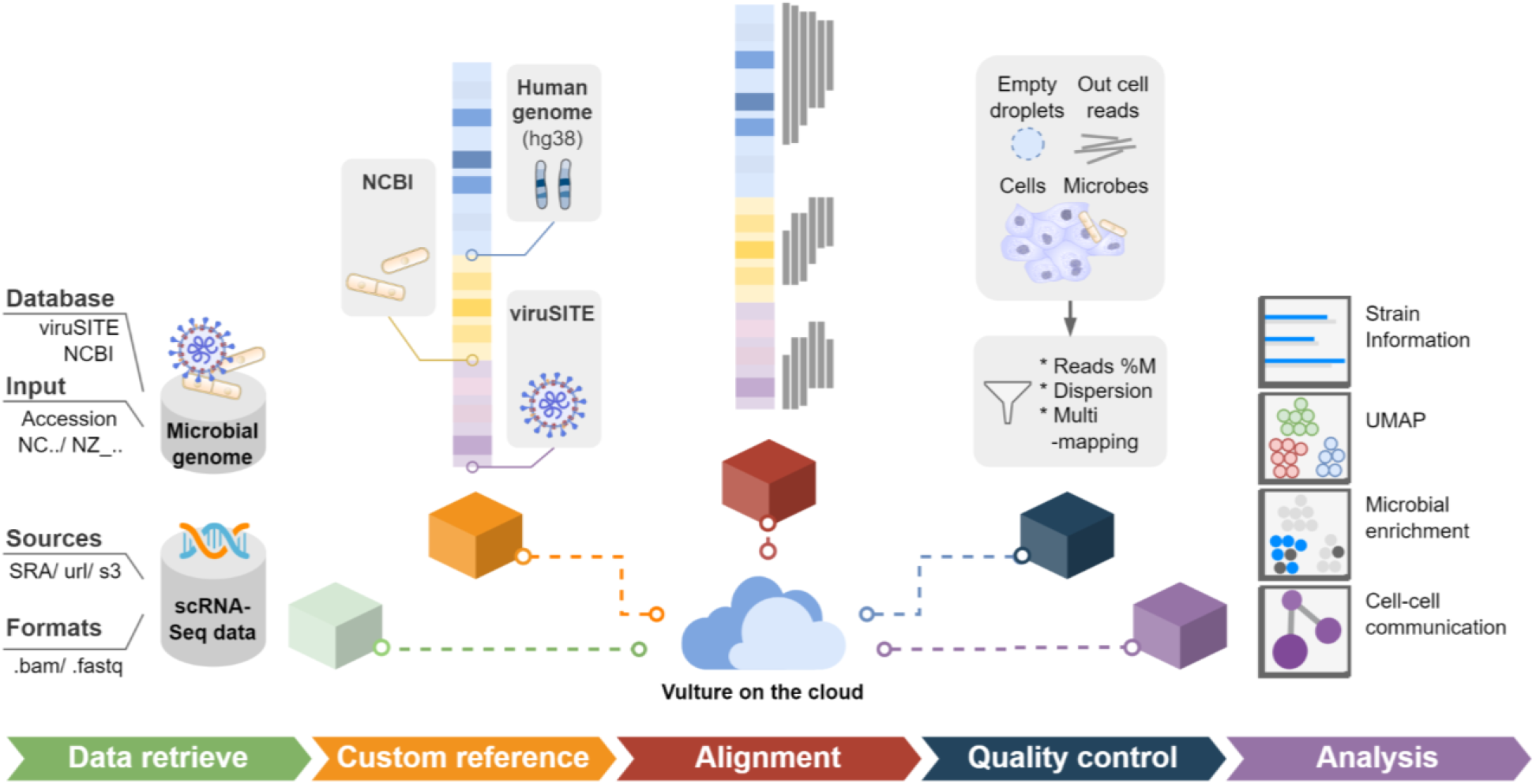
Schematic diagram of Vulture for scRNA-seq microbial calling on the cloud. Vulture is a containerized computational framework composed of five procedures. The five steps include 1) multi-format scRNA-seq data and microbial genome retrieval; 2) custom combined reference construction; 3) reads alignment; 4) quality control, and 5) downstream analysis. All procedures in the Vulture architecture run as containerized applications on the Amazon Batch service.

Vulture utilizes cloud computing to provide a fast, scalable, and cost-effective, viral calling framework without the need for hardware maintenance. Vulture is built on the AWS Batch service natively, which efficiently runs a huge amount of computing jobs while optimizing compute resources. At the same time, Vulture is implemented by a docker container and can be easily run on local servers or other cloud platforms. We applied Nextflow, a workflow management language to deploy complex parallel workflows of containers on clouds and clusters. The cloud architecture of Vulture is described in the Method section and **Supplementary Fig. S1b**. Through an AWS Batch and Nextflow, users can run thousands of viral calling tasks for public scRNA-seq data in parallel with simple configuration inputs.

### Runtime performance and cost-effectiveness of Vulture on the cloud

To validate and benchmark the scalability of Vulture on the cloud, we tested it through the public COVID-19 scRNA-seq data by Bost et al. [14] consisting of up to 400 run accessions. Execution duration (pastel) and vCPU time (saturated) from retrieving files to bam analysis of running 25 to 200 parallel tasks are recorded in **Fig. 2a**. Vulture analysis on 200 fastq files within 48 minutes, reaching a speed up (compared to running 200 single tasks sequentially) of 155x, showing that it is highly scalable. We also compared Vulture with another cloud-based method Cumulus [18]. Because Cumulus did not natively support viral calling tasks, we benchmark the read mapping speed and cost on the cloud only between two tools by aligning to an identical pre-built combined host-microbe genome. **Fig. 2b** indicates that Vulture outperformed Cumulus nearly twofold in total duration (saturated) though their per-task durations are close. Costing $12, Vulture runs 200 alignment tasks in 20 minutes, while Cumulus needs $69 to run 200 samples in 32 minutes. Vulture takes advantage of the AWS Batch spot capacity optimization technique. Its utilization of spot instances reduced the cost of running viral calling pipelines. Also, it ensured the availability of computational resource allocation, maximizing the number of concurrent tasks, and minimizing the response time of pending tasks.

**Fig. 2.**
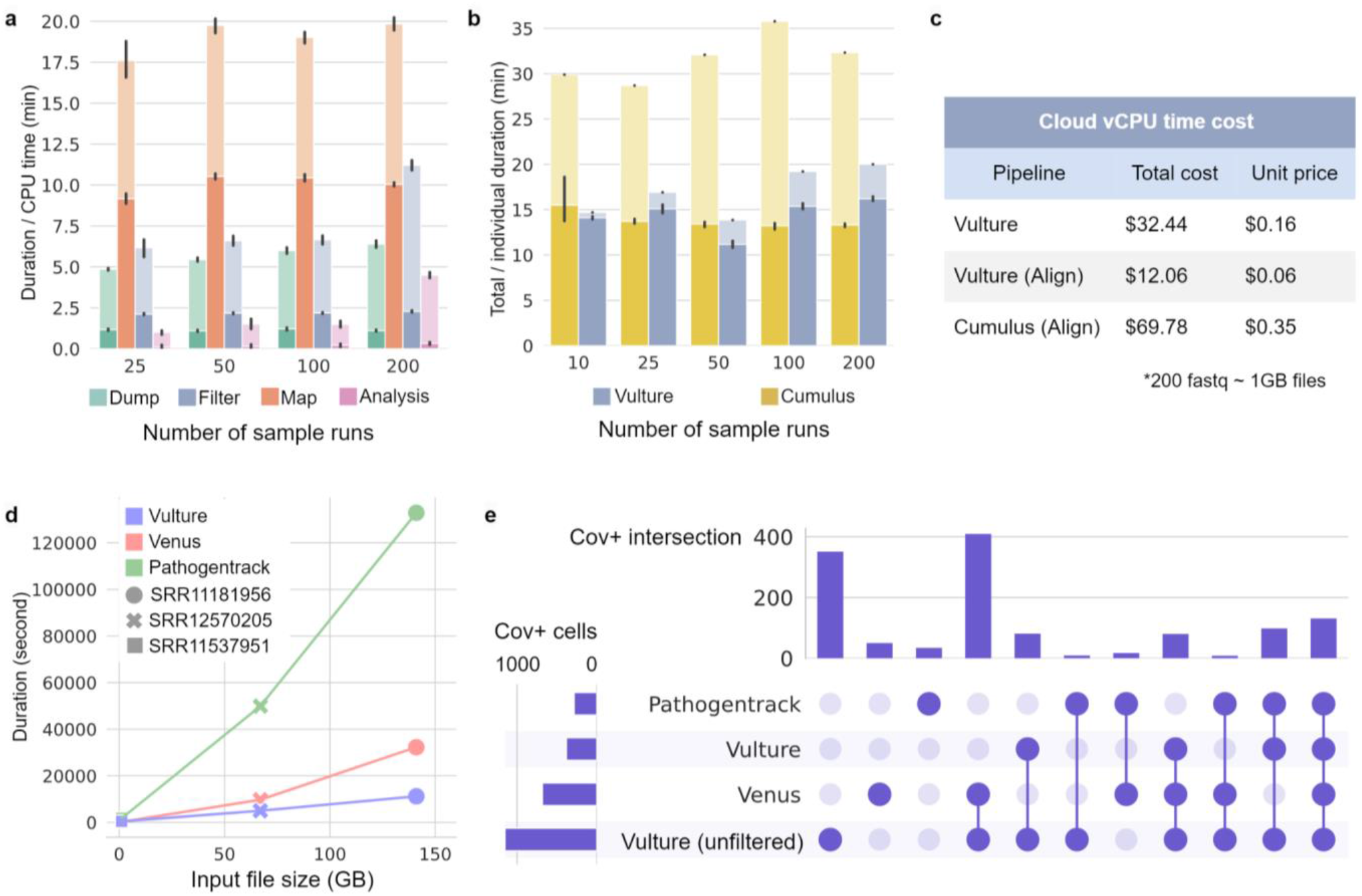
Performance benchmark of the Vulture. **a)** Performances of the Vulture pipeline to run 25 to 200 parallel analyses. The performance is measured by execution duration (pastel), and vCPU time (saturated) with the respective number of parallel tasks. The time of each of the four steps in the Vulture pipeline is displayed separately. **b)** Comparison between Vulture and Cumulus to run 25 to 200 parallel read alignment tasks. Wall clock time (pastel) and individual task durations (saturated) of different numbers parallel run processed by the two pipelines. **c)** Cost (in US dollars) of the computation resource needed to run 200 analyses on cloud platforms of Vulture and Cumulus. **d)** Task duration comparison among viral calling methods to run one single analysis with different input file sizes. **e)** Consistency among viral calling methods. The consistency is estimated by measuring the intersections of virus-positive cells annotated by different tools. Vulture results and results before the filtering step are discussed separately.

### Performance of Vulture on the local environment against off-the-shell tools

We also tested Vulture local command line tools against Venus and PathogenTrack [15], [16] using three COVID-19 scRNA-seq run with different file sizes: a 1GB small sample (SRR12570205) from Bost et al. [14], a 67 GB medium sample (SRR11537951) and a 141 GB large sample (SRR11181956) from Liao et al. [35]. **Fig. 2d** indicated that Vulture is the most computationally effective method among the three. In medium and large sample cases, Vulture took 5,009 and 11,225 seconds to finish one analysis. It was two to three-fold faster than Venus (9,734 and 32,282 seconds respectively) and nine to tenfolds faster than PathogenTrack (49,927 and 132,889 seconds respectively). Besides, we tested the consistency among three methods on three datasets by measuring the intersection of viral positive cell barcodes in **Fig. 2e**. The result of Vulture before filtering the empty droplets [28] named “Vulture (unfiltered)” is also added to the comparison. Since Vulture (unfiltered) is a superset of the Vulture result, the intersection between the two is the Vulture set in the figure. **Fig. 2e** indicates that Vulture reaches consensus with the others in most cases because most major sets are intersections with other methods. After all, the intersection of the three methods (which is usually the smallest) is the third largest in the test. Vulture’s result is particularly consistent with Venus with the largest set of intersections. Vulture (unfiltered) is the most sensitive with the largest number of SARS-CoV-2 positive cells (**Fig. 2e**) that cover most of the cells identified by Venus and PathogenTrack (**Supplementary Fig. S2a**). It is the quality control step that filtered out a large number of empty droplets. On 135 cells, the intersection of the three, we calculated the mean absolute error (MAE) and Pearson correlation of SARS-CoV-2 viral UMI counts across methods **Supplementary Fig. S2c and d**. The MAE between Vulture, Venus, and PathogenTrack are smaller than 1, showing that Vulture can consistently generate microbial calling results compared to state-of-the-art methods in a faster manner.

### Vulture enables cloud-based discovery of Metapneumovirus reads in COVID-19 BALF samples

SARS-CoV-2 infection has been identified to be the source of the worldwide COVID-19 pandemic since 2019. Many aspects of how the viral-host interaction have remained unrevealed. In particular, there is a major interest in identifying co-infection of other pathogens in COVID-19 patients. Therefore, we applied Vulture on the cloud on BALF samples from the Sequence Read Archive (SRA) to call viruses. We performed a meta-analysis on the Liao et al. [35] cohort from China (SRP250732) and the Bost et al. cohort [14] (SRP279746) from Israel. We ran a downstream analysis on BALF samples for two cohorts, totaling 51,338 and 991,722 cells after all QC filtering, respectively. After preprocessing and clustering the cell types were defined based on marker genes from Bost et al. and Liao et al. (**Supplementary Fig. S3b and c)**. Given the fact that SARS-CoV-2 UMIs in scRNA-seq data are relatively low and imbalanced, a statistical test (see methods) [19] is performed to estimate cell-type-specific enrichment of SAR-CoV-2 infection.

The combined microbe-host genome in Vulture includes a comprehensive set of human-host microbes to identify co-infections or unaware microbes. As a result, Vulture discovers the co-infection of the human metapneumovirus (hMPV) with SARS-CoV-2 in the Liao et al. [35] cohort (SRP250732), and unexpected Herpes simplex viruses (HSV) in the Bost et al. cohort [14] (SRP279746). This result is consistent with a previous discovery [14]. UMIs for different viral transcripts are shown in **Fig. 3a and b**. The visualization of BALF cells in the Liao cohort is shown in **Fig. 3c** and the presence of SARS-CoV-2 and hMPV is shown in **Fig. 3d and e**. UMAP plots for the Bost cohort and the presence of SARS-CoV-2 and HSV are shown in **Supplementary Fig. S3a.**The statistical test identified that SARS-CoV-2 is enriched (p-value < 0.05) in epithelial cells, neutrophils, and plasma B cells (**Fig. 3d and Table. 2**), and the infection of hMPV is enriched in CD8+ T cells, NK cells, macrophages, and monocytes (**Fig. 3e and Table. 4**). **Fig. 3e** demonstrates that a separated subtype of monocyte with numerous hMPV infections is presented. Therefore, we compare the differential expressed genes between the hMPV-enriched monocytes/macrophages to the hMPV-negative monocytes/macrophages. Differentially expressed genes for the viral infected subtype are shown in **Supplementary Table S1.** For instance, S100A8/S100A9 are up-regulated in the hMPV-enriched macrophages/monocytes, which are a heterodimer involved in neutrophil-related inflammatory processes [36]. FCN1 is also up-regulated which encodes a member of the complement cascade [3]. IDO1 is up-regulated, while infection with murine coronaviruses activates the AhR in an IDO1-independent manner, up-regulating the expression of proviral TCDD-inducible-PARP (TiPARP) and modulating cytokines [37]. We further applied g:Profiler [38] to search the functional enrichment of the top 100 up-regulated genes in hMPV-enriched subtypes. Results in **Fig. 3f** show that the expression pattern in the subtypes is highly responsible for the innate immune response, and cytokine responses. The interferon-gamma (IFN-γ) response which drives an IFN response in alveolar macrophages is reported to lead to the recruitment of monocyte-derived alveolar macrophages and then construct an inflammatory signaling circuit [36]. We also studied the cell-cell interaction (CCI) patterns of the SARS-CoV-2 and hMPV co-infection cases with the CellChat [33] tool. Each cell type in the CCI study is further divided into four subgroups including SARS-CoV-2 positive only (+-), hMPV positive only (-+), double positive (++), and normal cells (no markers). Virus-infected macrophages and hMPV-infected monocytes are tested to have stronger CCI than normal ones shown in **Fig. 3g**. Chemokine Signaling Pathways (CCL and CXCL) are tested to be the most significant signaling pathway (**Fig. 3h and i**) across the virally infected macrophages (+-/-+/++) and hMPV infected (-+) monocyte. Chemokine Signaling Pathways (CCL and CXCL) are reported to be involved in the pathogenesis of severe clinical sequelae of COVID-19, to SARS and MERS [39]. The top-ranked ligand-receptor (L-R) pairs among the CCL pathway are bindings of CCL3/5/7/8 to CCR1/5 **Supplementary Fig. S4**. Interaction of those L-R pairs induces monocyte recruitment into the lung parenchyma with subsequent differentiation into inflammatory macrophages and consecutive recruitment and activation of additional immune cells and epithelial damage [40]. The macrophage migration inhibitory factor (MIF) pathway, is described as a potential predictor for the outcome of critically ill patients and acute respiratory distress syndrome (ARDS) and a hallmark of severe COVID-19 disease [41]. We also observed a strong interaction of the GALECTIN signaling pathway among macrophages and monocytes. GALECTIN exhibits a pleiotropic role in mediating the acute and chronic consequences of infection and inflammation [42]. Other signaling pathways including ANNEXIN and SPP1 (**Fig3. h**) also contribute strong CCI among monocytes and macrophages.

**Fig. 3.**
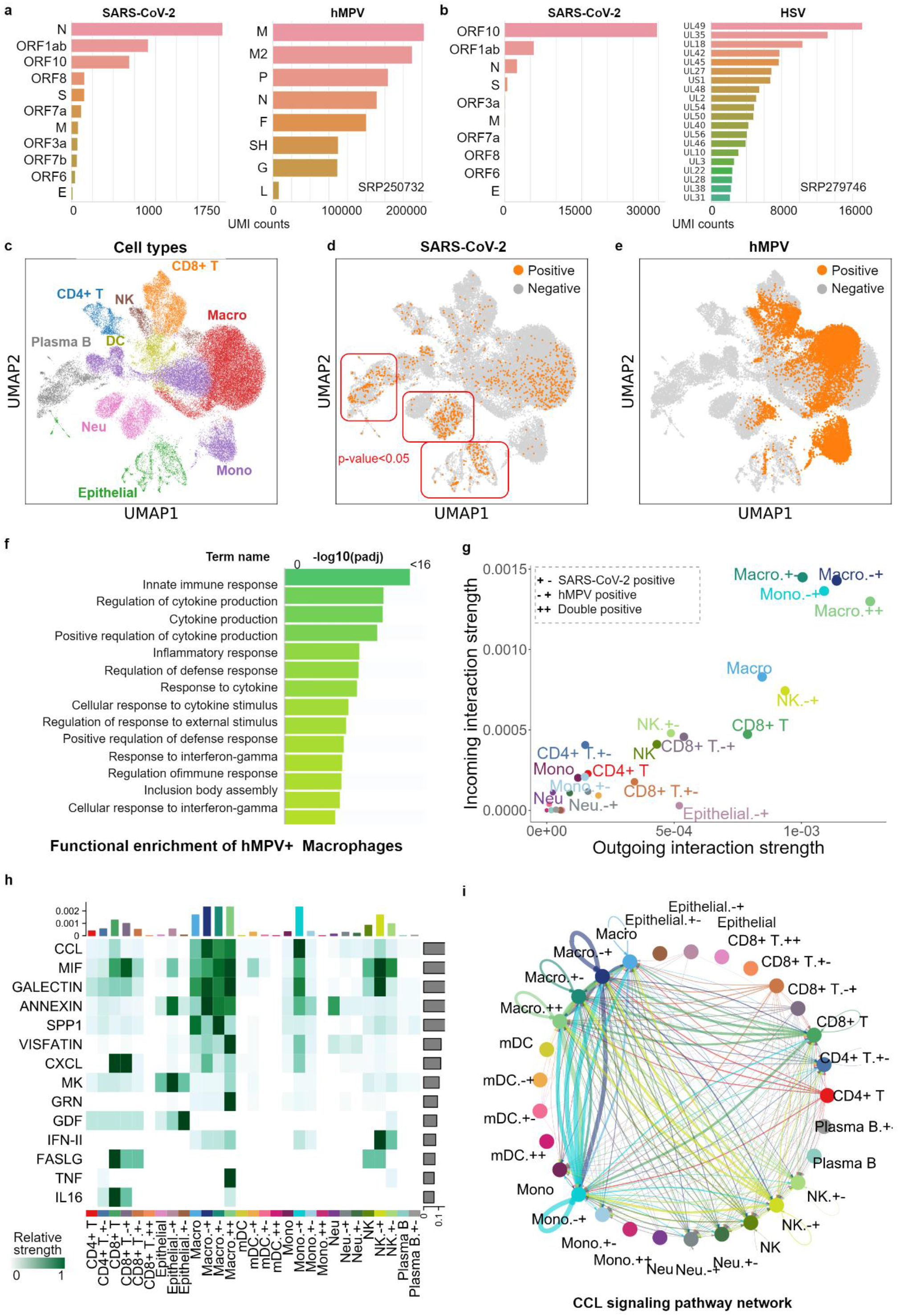
Viral calling meta-analysis results on the COVID-19 BALF samples. **a)** and **b)** are transcript UMIs of three major detected viruses (SARS-CoV-2, hMPV, and HSV) from SRP250732 and SRP 279746, respectively. **c)** UMAP plot of the COVID-19 BALF data, cells are colored by cell type annotations. **d)** and **e)** are UMAP plots of the SARS-CoV-2 and hMPV infection, respectively. Infected cells are colored orange while other cells are gray. **f)** is the top 15 enriched Gene Ontology (GO) terms identified by functional enrichment analysis. **g)** cell-cell interaction (CCI) strengths across different cells grouped by cell types along with viral infections. **h)** presents enriched signaling pathways identified by CellChat. **i)** is the CCI among cell types through the CCL signaling pathway network.

**Table. 1.** Overview of the datasets processed in this study.

| Data name | SRP accession | Disease | Tissue | #of runs | #of patients | Format |
| --- | --- | --- | --- | --- | --- | --- |
| Liao et al. [35] | SRP250732 | COVID-19 | BALF | 12 | 12 | fastq |
| Bost et al. [14] | SRP279746 | COVID-19 | BALF | 336 | 22 | fastq |
| Losic et al. [8] | SRP136347 | HCC | Tumor | 7 | 2 | bam |
| Sharma et al. [9] | SRP278381 | HCC | Tumor | 58 | 16 | bam |
| Ho et al. [10] | SRP318499 | HCC | Tumor | 8 | 8 | bam |
| Zhang et al. [11] | SRP215370 | GC | Tumor | 16 | 2 | fastq |
| Kim et al. [12] | SRP261119 | GC | Tumor | 13 | 13 | fastq |

**Table. 2.** Statistic of SARS-CoV-2 infection of cells in the BALF cohort. Cell types with bold text are enriched cell types.

| Cell types | Enrichment score | Proportion | Adjusted p-value | p-value |
| --- | --- | --- | --- | --- |
| **SRP250732** |  |  |  |  |
| CD4+ T | 0.003083 | 1.93% | 1 | 0.7686 |
| CD8+ T | 0.003053 | 0.65% | 1 | 1 |
| **Epithelial** | **0.004403** | **6.84%** | **<0.0001** | **<0.0001** |
| Macro | 0.005222 | 1.76% | 1 | 1 |
| Mono | 0.004527 | 1.27% | 1 | 1 |
| NK | 0.002431 | 1.03% | 1 | 0.9996 |
| **Neu** | **0.004833** | **5.20%** | **<0.0001** | **<0.0001** |
| **Plasma B** | **0.004589** | **4.59%** | **<0.0001** | **<0.0001** |
| DC | 0.002991 | 1.09% | 1 | 1 |
| **SRP279746** |  |  |  |  |
| Epithelial | 0.004675 | 0.12% | 1 | 0.4046 |
| Macro | 0.003758 | 0.13% | 1 | 0.9992 |
| Mono | 0.00313 | 0.15% | 1 | 1 |
| NK | 0.002682 | 0.13% | 1 | 1 |
| **Neu** | **0.005471** | **0.19%** | **<0.0001** | **<0.0001** |
| Plasma B | 0 | 0.00% | 1 | 1 |

**Table. 3.**
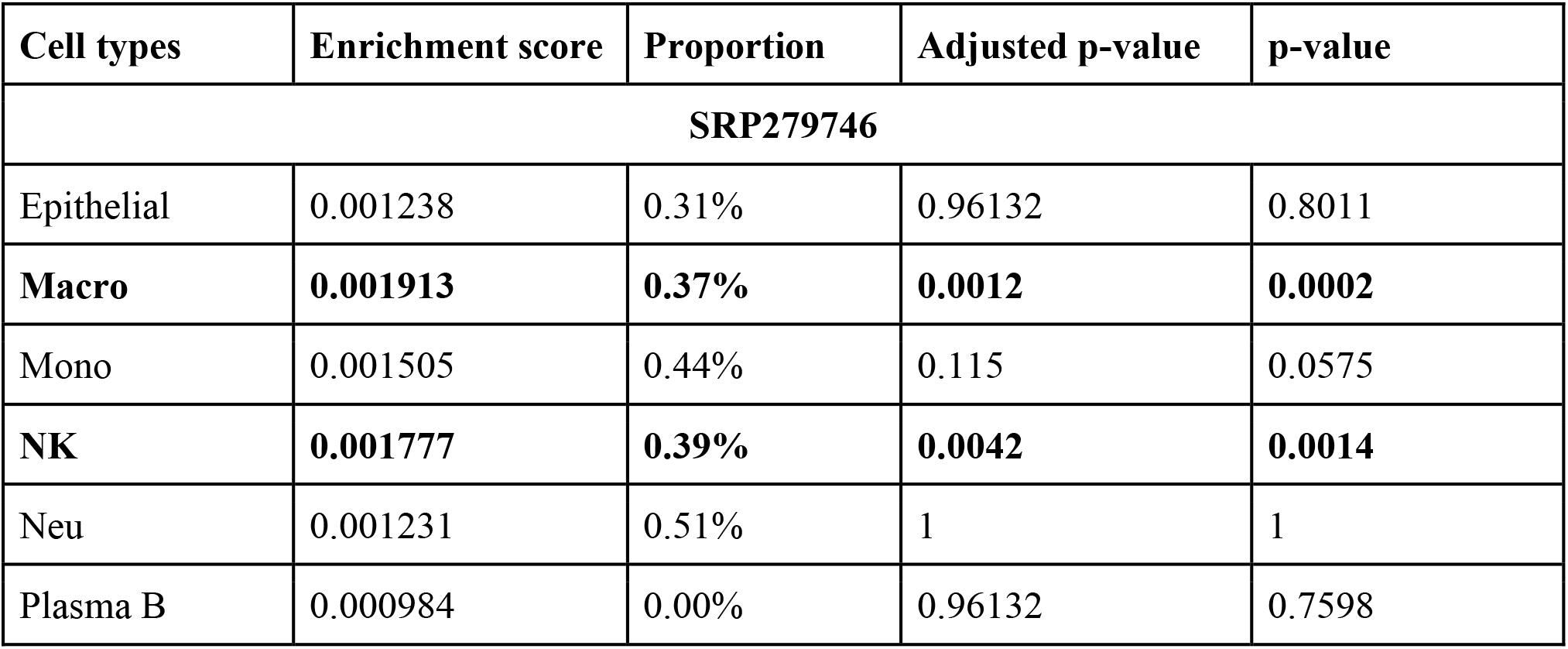
Statistic of intracellular HSV in the Bost et al. cohort. Cell types with bold text are enriched cell types.

| Cell types | Enrichment score | Proportion | Adjusted p-value | p-value |
| --- | --- | --- | --- | --- |
| **SRP279746** |  |  |  |  |
| Epithelial | 0.001238 | 0.31% | 0.96132 | 0.8011 |
| **Macro** | **0.001913** | **0.37%** | **0.0012** | **0.0002** |
| Mono | 0.001505 | 0.44% | 0.115 | 0.0575 |
| **NK** | **0.001777** | **0.39%** | **0.0042** | **0.0014** |
| Neu | 0.001231 | 0.51% | 1 | 1 |
| Plasma B | 0.000984 | 0.00% | 0.96132 | 0.7598 |

**Table. 4.** Statistic of intracellular hMPV in the Liao et al. cohort. Cell types with bold text are enriched cell types.

| Cell types | Enrichment score | Proportion | Adjusted p-value | p-value |
| --- | --- | --- | --- | --- |
| **SRP250732** |  |  |  |  |
| CD4+ T | -0.003013 | 0.00% | 1 | 1 |
| **CD8+ T** | **0.000598** | **27.68%** | **<0.0001** | **<0.0001** |
| Epithelial | 0.000396 | 5.78% | 1 | 1 |
| **Marco** | **0.00073** | **32.23%** | **<0.0001** | **<0.0001** |
| **Mono** | **0.000677** | **26.95%** | **<0.0001** | **<0.0001** |
| **NK** | **0.000535** | **41.27%** | **<0.0001** | **<0.0001** |
| Neu | 0.000475 | 6.97% | 1 | 1 |
| Plasma B | -0.003013 | 0.00% | 1 | 1 |
| mDC | 0.000454 | 8.90% | 1 | 1 |

### Cloud-based meta-analysis reveals an HBV-associated CNV signature in HCC

Another advantage of having a cloud-based framework is that it facilitates the integration of multiple data sets that are already in the same repository on the cloud. We performed a meta-analysis of the three public hepatocellular carcinoma (HCC) cohorts with droplet scRNA-seq sequencing data, SRP278381, SRP136347, and SRP318499. We ran Vulture on the HCC samples of 24 patients in these cohorts, totaling 421,780 cells following all QC filtering. After clustering and integration, the cell types were defined based on marker genes from Sharma et al. (SRP278381) (**Supplementary Fig. S5c**). Microbial enrichment detection (**Table. 5**) on each of the cohorts indicated hepatocytes are the only cell type with HBV enrichment (**Supplementary Fig. S5b**).

**Table. 5.** Statistics of intracellular HBV of cells in the HCC cohort. Cell types with bold text are enriched cell types.

| Cell types | Enrichment score | Proportion | Adjusted p-value | p-value |
| --- | --- | --- | --- | --- |
| **SRP136347** |  |  |  |  |
| B cells | 0.001301 | 0.46% | 1 | 0.9958 |
| Endothelial | 0.002117 | 0.64% | 1 | 0.9954 |
| Fibroblast | 0.001942 | 0.64% | 1 | 0.9896 |
| **Hepatocyte** | **0.003583** | **0.98%** | **<0.0001** | **<0.0001** |
| Myeloid | 0.002117 | 0.43% | 1 | 1 |
| T cells | 0.002249 | 0.41% | 1 | 1 |
| **SRP278381** |  |  |  |  |
| B cells | 1.38E-19 | 0.02% | 1 | 1 |
| Endothelial | 0.00245 | 0.27% | 1 | 1 |
| Fibroblast | 0.001206 | 0.16% | 1 | 1 |
| **Hepatocyte** | **0.004524** | **3.24%** | **<0.0001** | **<0.0001** |
| Myeloid | 0.002109 | 0.29% | 1 | 1 |
| T cells | 0.002374 | 0.10% | 1 | 1 |
| **SRP318499** |  |  |  |  |
| B cells | 0.000821 | 4.22% | 1 | 1 |
| Endothelial | 0.000839 | 3.90% | 1 | 1 |
| Fibroblast | 0.000579 | 2.74% | 1 | 1 |
| **Hepatocyte** | **0.001857** | **15.33%** | **<0.0001** | **<0.0001** |
| Myeloid | 0.001243 | 4.77% | 1 | 1 |
| T cells | 0.001334 | 3.51% | 1 | 1 |

To further delineate the HBV enrichment within hepatocytes, we re-clustered and re-integrated the hepatocytes of 11/24 patient samples that contained any HBV expression (**Fig. 4a**), and found that the only HBV-enriched subclusters were 0 and 3 (**Table. 6**), with both subclusters enriched in 2/3 cohorts (**Fig. 4b**). We analyzed the CNV of each patient using inferCNV with the macrophages as reference cells and hepatocytes as observation cells (**Supplementary Fig. S6-8**); we noted generally the CNV clones with more HBV expression indeed mostly consisted of subclusters 0 and 3, while the clones with less HBV expression consisted of mainly subcluster 2. We picked three representative patients (from two cohorts) with well-defined CNV clones and a sufficient (>50) number of HBV-positive cells, P114_SRP318499, P725_SRP318499, P7_SRP278381, and generated an overall CNV for this set (**Fig. 4c**). The result shows that for patients in different cohorts, the clones with discernable, less ambiguous CNV patterns have a clear majority of cells with HBV expression compared to the other clones (**Fig. 4c**, green boxes).

**Fig. 4.**
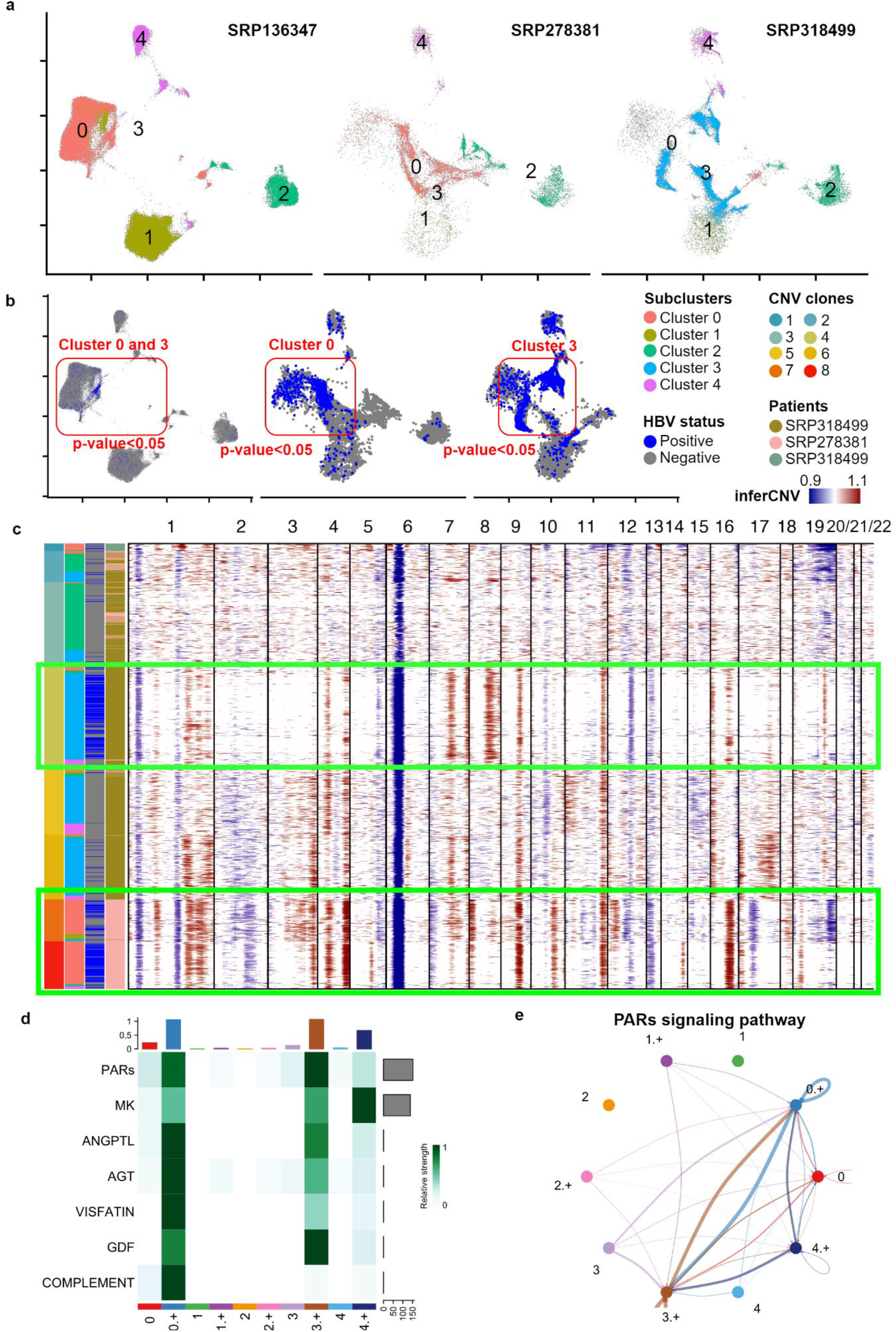
Viral calling meta-analysis results on the HCC hepatocytes. **a)** is a UMAP plot of the HBV-enriched hepatocytes, cells are colored by sub-clustering annotations. **b)** are UMAP plots of the intracellular HBV. HBV-positive cells are colored blue while other cells are gray. **c)** is the top 15 enriched Gene Ontology (GO) terms identified by functional enrichment analysis. **c)** heatmap of CNV inferred by inferCNV across chromosomes on the HCC hepatocytes. Cells are labeled by different annotations including CNV clones, sub-clusters, HBV status, and patients. **d)** presents enriched signaling pathways identified by CellChat of hepatocytes. **e)** is the CCI among cell types through the PARs signaling pathway network.

**Table. 6.** Statistics of intracellular HBV of hepatocyte cells in the HCC cohort. Cell types with bold text are enriched cell types.

| Hepatocyte subtypes | Enrichment score | Proportion | Adjusted p-value | p-value |
| --- | --- | --- | --- | --- |
| **SRP136347** |  |  |  |  |
| 0 | 0.001419 | 17.46% | 0.1183 | 0.0338 |
| 1 | 0.000997 | 3.41% | 1 | 1 |
| 2 | 0.00122 | 2.95% | 1 | 1 |
| **3** | **0.002167** | **19.48%** | **0** | **0** |
| 4 | 0.00142 | 9.28% | 1 | 1 |
| **SRP278381** |  |  |  |  |
| **0** | **0.00488** | **21.01%** | **<0.0001** | **<0.0001** |
| 1 | 0.002635 | 7.27% | 1 | 1 |
| 2 | 0.001877 | 0.46% | 1 | 1 |
| 3 | 0.001957 | 6.72% | 1 | 0.9992 |
| 4 | 0.002577 | 5.18% | 1 | 1 |
| **SRP318499** |  |  |  |  |
| **0** | **0.004107** | **3.28%** | **<0.0001** | **<0.0001** |
| 1 | 0.003232 | 0.64% | 1 | 1 |
| 2 | 0.002589 | 0.73% | 1 | 1 |
| **3** | **0.000926** | **13.89%** | **0.0003** | **0.0001** |
| 4 | 0.002699 | 1.20% | 1 | 1 |

We studied the CCI pattern of HBV-hepatocyte interactions and further grouped hepatocytes into HBV-positive (+) and normal (not HBV-positive) hepatocytes. CCIs in HBV-positive subclusters 0 and 3 have higher relative strength than normal ones shown in **Fig. 4d**. Proteinase-activated receptors (PARs) signaling pathway is tested to be the most significant signaling pathway (**Fig. 4d and e**) across the HBV-enriched hepatocytes. PARs as the thrombin receptors are involved in thrombin-induced cell migration across a collagen transmembrane barrier [43]. Midkine (MK) is a growth factor that is tested to be a crucial role in HCC. It is involved in inflammatory responses, acts as an anti-apoptotic factor, and blocks anoikis to promote metastasis [44].

### Identification of *H. pylori* reads in gastric cancer

We also conducted a meta-analysis of the two gastric cancer (GC) cohorts that have publicly available droplet scRNA-seq sequencing data, SRP215370 (Zhang et al. [11]) and SRP261119 (Kim et al. [12]). The former includes early GC samples (which we only used the two confirmed *H. pylori*+ patients) and the latter contains GC patient samples, the majority of which were known to be *H. pylori*+. In total, we applied Vulture on the early GC or GC samples of 15 patients in the two cohorts, amounting to 125,845 after all QC filtering. We used the same approach as the above HCC case study for clustering and integration and labeled the cell types in line with Kim et al. (SRP261119) (**Fig. 5a**). Microbial enrichment detection by cohort showed *H. pylori* enrichment in endothelial cells, fibroblasts, and macrophages in SRP215370, and enrichment only in pit mucous cells in SRP261119 (**Fig. 5c** and **Table. 7**). The difference is also present in the *H. pylori* virulence of two cohorts shown in **Fig. 3b**. CagA virulence gene is only detected in the Zhang et al. cohort where cagA-positive strains are the strongest risk factor of gastric cancer [45].

**Fig. 5.**
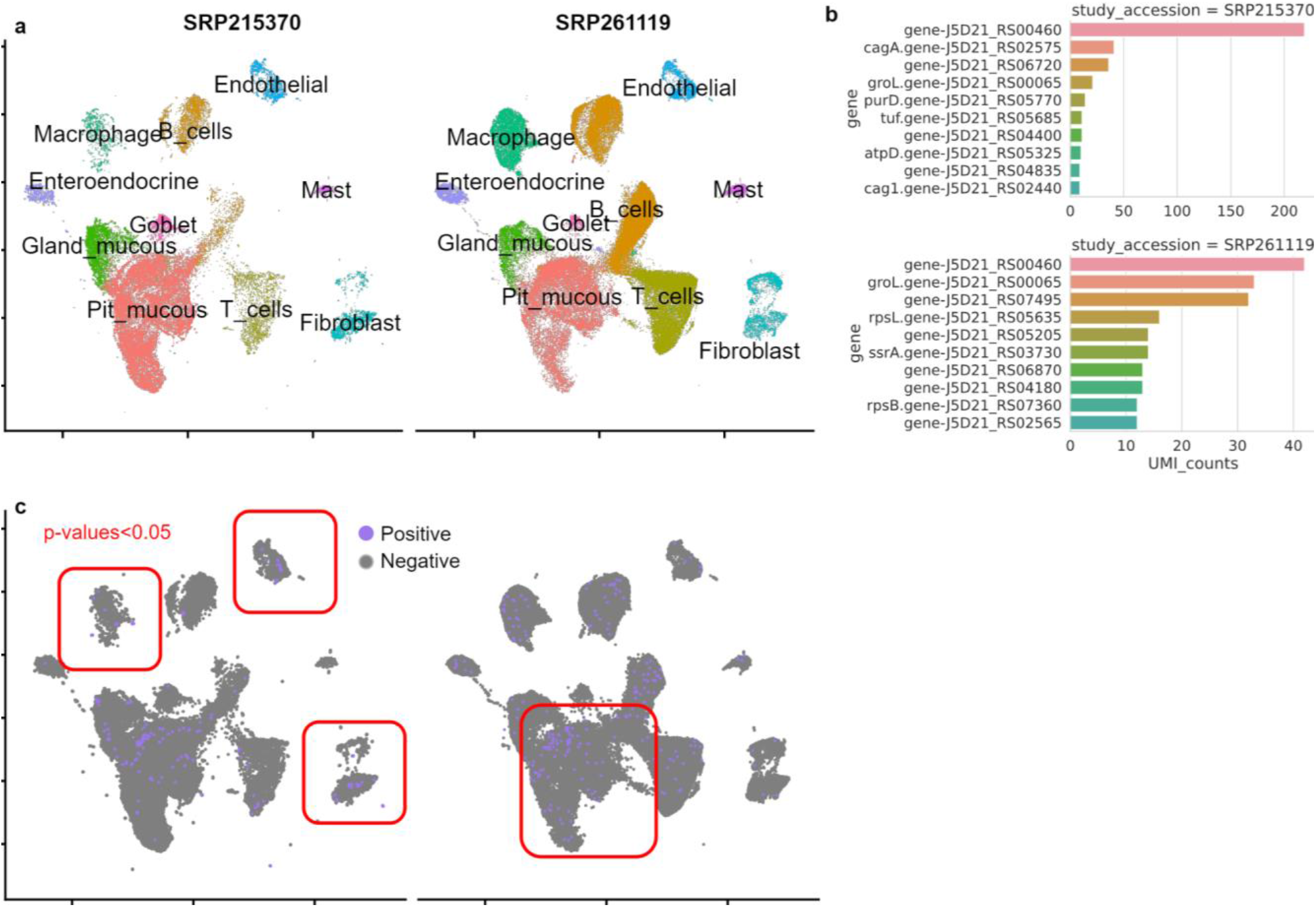
Viral calling meta-analysis results on the GC sample. **a)** is a UMAP plot of the GC cells colored by cell type annotations. **b)** is the transcript UMIs of *H. pylori* identified in SRP215370 and SRP2161119, separately. **c)** are UMAP plots of the intracellular *H. pylori. H. pylori-positive* cells are colored purple while other cells are gray.

**Table. 7.** Statistics of intracellular *H. pylori* of cells in the GC cohort. Cell types with bold text are enriched cell types.

| Cell types | Enrichment score | Proportion | Adjusted p-value | p-value |
| --- | --- | --- | --- | --- |
| **SRP215370** |  |  |  |  |
| B cells | 0.012183 | 0.28% | 1 | 0.932 |
| **Endothelial** | **0.014991** | **0.99%** | **0.003** | **0.0009** |
| Enteroendocrine | 1.17E-18 | 0.10% | 1 | 0.984 |
| **Fibroblast** | **0.019649** | **1.54%** | **<0.0001** | **<0.0001** |
| Gland_mucous | 0.014328 | 0.26% | 1 | 0.9848 |
| Goblet | 1.17E-18 | 0.07% | 1 | 0.9988 |
| **Macrophage** | **0.016109** | **1.92%** | **<0.0001** | **<0.0001** |
| Mast | -0.19071 | 0.00% | 1 | 1 |
| Pit mucous | 0.022554 | 0.28% | 1 | 1 |
| T cells | 0.013148 | 0.52% | 0.642 | 0.2568 |
| **SRP261119** |  |  |  |  |
| B cells | 0.018981 | 0.21% | 1 | 0.9872 |
| Endothelial | 0.007556 | 0.24% | 1 | 0.6366 |
| Enteroendocrine | 0.010316 | 0.52% | 0.215 | 0.043 |
| Fibroblast | 0.007556 | 0.15% | 1 | 0.9507 |
| Gland_mucous | 0.009763 | 0.21% | 1 | 0.7916 |
| Goblet | -0.16922 | 0.00% | 1 | 1 |
| Macrophage | 0.014721 | 0.32% | 0.695 | 0.2085 |
| Mast | 0.006508 | 0.24% | 1 | 0.656 |
| **Pit mucous** | **0.020573** | **0.47%** | **<0.0001** | **<0.0001** |
| T cells | 0.014512 | 0.14% | 1 | 1 |

## Discussion

In recent years, the rapidly growing single-cell study sources have become a gold mine for the re-investigation of host-microbial interactions. But neither traditional practices on scRNA-seq studies nor the bioinformatics methods developed to detect microbes are capable of scalable meta-analysis for the public open data on the cloud. Alternatively, cloud computing has become an essential piece of equipment. Here, we present Vulture, a cloud-based scalable framework for calling microbial RNA on public scRNA-seq resources. Vulture was benchmarked on data originating from various tissues, generated with different scRNA-seq platforms, and deposited in different formats. It is tested to be highly scalable and cost-effective. Because it runs 200 analyses within a similar duration and low cost compared to a single task. Moreover, running a single Vulture analysis on the local environment is significantly faster than previous methods. The reason is that Vulture provides an easily customizable combined reference. It performs read mapping once on while others map read to the host and then align unmapped reads to the microbe references, among other intermediate steps. Vulture runs faster by getting rid of complex preprocessing on unmapped reads. The combined reference is 30% larger than the host genome but indexing from alignment tools can compensate for the increased reference size. Also, Vulture is user-friendly because it supports multiple platforms and multiple input formats. We demonstrated that Vulture can readily provide an effective solution for viral calling meta-analysis on large-scale public data.

We applied Vulture and scRNA-seq analysis to public COVID-19 BALF cells, HCC samples, and GC samples. The COVID-19 analysis revealed that Vulture is capable of identifying co-infections of unexpected pathogens. The HCC samples discovered a potential crucial relationship between CNV and intracellular HBV. All those cases indicate that Vulture is highly valuable to study unknown mechanisms and treatments by mining large-scale single-cell data.

However, there are several limitations of Vulture. A key hinder to Vulture meta-analysis is the permission of data. Serval atlas-level databases have strict access permission which makes it hard to run cloud-based analysis on a large scale. Also, large-scale meta-data cleansing for raw sequencing files, which is essential to run parallel meta-analysis appropriately is difficult because sequencing files are generated by different protocols. A prospective solution is to incorporate biomedical natural language processing (bioNLP) models and search engine technologies to subtract metadata for Vulture. Ultimately, Vulture automatically digs the gold mine of host-microbial interactions in big data.

In summary, Vulture is a cloud-based scalable framework for calling microbial RNA on public scRNA-seq resources. It is not only highly scalable, cost-effective on the cloud, and significantly outperformed previous methods in local environments. We anticipate that Vulture will play a crucial role in the attempt to understand the unrevealed genetics of pathogenic diseases as the community gradually contributes to the increasing scale of single-cell data for host-microbial interactions

## Data Availability

Source code and tutorial of Vulture is deposited on GitHub (https://github.com/holab-hku/Vulture). The datasets supporting the conclusions of this article are available in the SRA Run Selector (https://www.ncbi.nlm.nih.gov/Traces/study/) repository. They can be searched under the following project accessions numbers: SRP250732 (Liao et al. [35]), SRP279746 (Bost et al. [14]), SRP136347 (Losic et al. [8]), SRP278381 (Sharma et al. [9]), SRP318499 (Ho et al. [10]), SRP215370 (Zhang et al. [11]), and SRP261119 (Kim et al. [12]).

## Acknowledgements

JWKH conceived and designed the study. JC, KHOY, and DY designed and implemented the computational pipeline and the cloud framework. HYHW contributed to the collection of data. XD contributes to the testing of prototypes. JC and KHOY performed the case study analysis and implemented the data analytics. All authors wrote, read, reviewed the manuscript, and approved the final version.

The authors thank all their colleagues, particularly at D^2^4H and The University of Hong Kong for their support and intellectual engagement.

## References

[1] M. Levrero and J. Zucman-Rossi, “Mechanisms of HBV-induced hepatocellular carcinoma,” J. Hepatol., vol. 64, no. 1, Supplement, pp. S84–S101, Apr. 2016, doi: 10.1016/j.jhep.2016.02.021.

[2] L. E. Wroblewski, R. M. Peek, and K. T. Wilson, “Helicobacter pylori and Gastric Cancer: Factors That Modulate Disease Risk,” Clin. Microbiol. Rev., vol. 23, no. 4, pp. 713–739, Oct. 2010, doi: 10.1128/CMR.00011-10.

[3] Y. Tian, L. N. Carpp, H. E. R. Miller, M. Zager, E. W. Newell, and R. Gottardo, “Single-cell immunology of SARS-CoV-2 infection,” Nat. Biotechnol., vol. 40, no. 1, Art. no. 1, Jan. 2022, doi: 10.1038/s41587-021-01131-y.

[4] N. Drayman, P. Patel, L. Vistain, and S. Tay, “HSV-1 single-cell analysis reveals the activation of anti-viral and developmental programs in distinct sub-populations,” eLife, vol. 8, p. e46339, May 2019, doi: 10.7554/eLife.46339.

[5] M. Shnayder et al., “Defining the Transcriptional Landscape during Cytomegalovirus Latency with Single-Cell RNA Sequencing,” mBio, vol. 9, no. 2, pp. e00013–18, Mar. 2018, doi: 10.1128/mBio.00013-18.

[6] Y. Steuerman et al., “Dissection of Influenza Infection In Vivo by Single-Cell RNA Sequencing,” Cell Syst., vol. 6, no. 6, pp. 679–691.e4, Jun. 2018, doi: 10.1016/j.cels.2018.05.008.

[7] F. Zanini et al., “Virus-inclusive single-cell RNA sequencing reveals the molecular signature of progression to severe dengue,” Proc. Natl. Acad. Sci., vol. 115, no. 52, Dec. 2018, doi: 10.1073/pnas.1813819115.

[8] B. Losic et al., “Intratumoral heterogeneity and clonal evolution in liver cancer,” Nat. Commun., vol. 11, no. 1, Art. no. 1, Jan. 2020, doi: 10.1038/s41467-019-14050-z.

[9] A. Sharma et al., “Onco-fetal Reprogramming of Endothelial Cells Drives Immunosuppressive Macrophages in Hepatocellular Carcinoma,” Cell, vol. 183, no. 2, pp. 377–394.e21, Oct. 2020, doi: 10.1016/j.cell.2020.08.040.

[10] D. W.-H. Ho et al., “Single-cell RNA sequencing shows the immunosuppressive landscape and tumor heterogeneity of HBV-associated hepatocellular carcinoma,” Nat. Commun., vol. 12, no. 1, Art. no. 1, Jun. 2021, doi: 10.1038/s41467-021-24010-1.

[11] P. Zhang et al., “Dissecting the Single-Cell Transcriptome Network Underlying Gastric Premalignant Lesions and Early Gastric Cancer,” Cell Rep., vol. 27, no. 6, pp. 1934–1947.e5, May 2019, doi: 10.1016/j.celrep.2019.04.052.

[12] J. Kim et al., “Single-cell analysis of gastric pre-cancerous and cancer lesions reveals cell lineage diversity and intratumoral heterogeneity,” Npj Precis. Oncol., vol. 6, no. 1, Art. no. 1, Jan. 2022, doi: 10.1038/s41698-022-00251-1.

[13] A. Regev et al., “The Human Cell Atlas,” eLife, vol. 6, p. e27041, Dec. 2017, doi: 10.7554/eLife.27041.

[14] P. Bost et al., “Host-Viral Infection Maps Reveal Signatures of Severe COVID-19 Patients,” Cell, vol. 181, no. 7, pp. 1475–1488.e12, Jun. 2020, doi: 10.1016/j.cell.2020.05.006.

[15] W. Zhang, X. Xu, Z. Fu, J. Chen, S. Chen, and Y. Tan, “PathogenTrack and Yeskit: tools for identifying intracellular pathogens from single-cell RNA-sequencing datasets as illustrated by application to COVID-19,” Front. Med., vol. 16, no. 2, pp. 251–262, Apr. 2022, doi: 10.1007/s11684-021-0915-9.

[16] C. Y. Lee et al., “Venus: An efficient virus infection detection and fusion site discovery method using single-cell and bulk RNA-seq data,” PLOS Comput. Biol., vol. 18, no. 10, p. e1010636, Oct. 2022, doi: 10.1371/journal.pcbi.1010636.

[17] A. Yang, M. Troup, P. Lin, and J. W. Ho, “Falco: a quick and flexible single-cell RNA-seq processing framework on the cloud,” Bioinformatics, vol. 33, no. 5, pp. 767–769, 2017.

[18] B. Li et al., “Cumulus provides cloud-based data analysis for large-scale single-cell and single-nucleus RNA-seq,” Nat. Methods, vol. 17, no. 8, pp. 793–798, 2020.

[19] T. M. Delorey et al., “COVID-19 tissue atlases reveal SARS-CoV-2 pathology and cellular targets,” Nature, vol. 595, no. 7865, pp. 107–113, 2021.

[20] R. C. Edgar et al., “Petabase-scale sequence alignment catalyses viral discovery,” Nature, pp. 1–6, 2022.

[21] D. Karolchik et al., “The UCSC genome browser database,” Nucleic Acids Res., vol. 31, no. 1, pp. 51–54, 2003.

[22] M. Stano, G. Beke, and L. Klucar, “viruSITE—integrated database for viral genomics,” Database, vol. 2016, 2016.

[23] H. Li, “Minimap2: pairwise alignment for nucleotide sequences,” Bioinformatics, vol. 34, no. 18, pp. 3094–3100, Sep. 2018, doi: 10.1093/bioinformatics/bty191.

[24] B. Kaminow, D. Yunusov, and A. Dobin, “STARsolo: accurate, fast and versatile mapping/quantification of single-cell and single-nucleus RNA-seq data,” bioRxiv, 2021.

[25] G. X. Y. Zheng et al., “Massively parallel digital transcriptional profiling of single cells,” Nat. Commun., vol. 8, no. 1, p. 14049, Jan. 2017, doi: 10.1038/ncomms14049.

[26] P. Melsted et al., “Modular, efficient and constant-memory single-cell RNA-seq preprocessing,” Nat. Biotechnol., vol. 39, no. 7, pp. 813–818, 2021.

[27] A. Srivastava, L. Malik, T. Smith, I. Sudbery, and R. Patro, “Alevin efficiently estimates accurate gene abundances from dscRNA-seq data,” Genome Biol., vol. 20, no. 1, pp. 1–16, 2019.

[28] A. T. Lun, S. Riesenfeld, T. Andrews, T. Gomes, J. C. Marioni, and others, “EmptyDrops: distinguishing cells from empty droplets in droplet-based single-cell RNA sequencing data,” Genome Biol., vol. 20, no. 1, pp. 1–9, 2019.

[29] F. A. Wolf, P. Angerer, and F. J. Theis, “SCANPY: large-scale single-cell gene expression data analysis,” Genome Biol., vol. 19, no. 1, pp. 1–5, 2018.

[30] R. Satija, J. A. Farrell, D. Gennert, A. F. Schier, and A. Regev, “Spatial reconstruction of single-cell gene expression data,” Nat. Biotechnol., vol. 33, no. 5, pp. 495–502, 2015.

[31] K. Polański, M. D. Young, Z. Miao, K. B. Meyer, S. A. Teichmann, and J.-E. Park, “BBKNN: fast batch alignment of single cell transcriptomes,” Bioinformatics, vol. 36, no. 3, pp. 964–965, Feb. 2020, doi: 10.1093/bioinformatics/btz625.

[32] I. Korsunsky et al., “Fast, sensitive and accurate integration of single-cell data with Harmony,” Nat. Methods, vol. 16, no. 12, Art. no. 12, Dec. 2019, doi: 10.1038/s41592-019-0619-0.

[33] S. Jin et al., “Inference and analysis of cell-cell communication using CellChat,” Nat. Commun., vol. 12, no. 1, pp. 1–20, 2021.

[34] A. P. Patel et al., “Single-cell RNA-seq highlights intratumoral heterogeneity in primary glioblastoma,” Science, vol. 344, no. 6190, pp. 1396–1401, 2014.

[35] M. Liao et al., “Single-cell landscape of bronchoalveolar immune cells in patients with COVID-19,” Nat. Med., vol. 26, no. 6, pp. 842–844, 2020.

[36] M. Mahler, P.-L. Meroni, M. Infantino, K. A. Buhler, and M. J. Fritzler, “Circulating Calprotectin as a Biomarker of COVID-19 Severity,” Expert Rev. Clin. Immunol., vol. 17, no. 5, pp. 431–443, May 2021, doi: 10.1080/1744666X.2021.1905526.

[37] W. A. Turski, A. Wnorowski, G. N. Turski, C. A. Turski, and L. Turski, “AhR and IDO1 in pathogenesis of Covid-19 and the ‘Systemic AhR Activation Syndrome:’ a translational review and therapeutic perspectives,” Restor. Neurol. Neurosci., vol. 38, no. 4, pp. 343–354, Sep. 2020, doi: 10.3233/RNN-201042.

[38] U. Raudvere et al., “g:Profiler: a web server for functional enrichment analysis and conversions of gene lists (2019 update),” Nucleic Acids Res., vol. 47, no. W1, pp. W191–W198, Jul. 2019, doi: 10.1093/nar/gkz369.

[39] F. Coperchini, L. Chiovato, L. Croce, F. Magri, and M. Rotondi, “The cytokine storm in COVID-19: An overview of the involvement of the chemokine/chemokine-receptor system,” Cytokine Growth Factor Rev., vol. 53, pp. 25–32, Jun. 2020, doi: 10.1016/j.cytogfr.2020.05.003.

[40] R. L. Chua et al., “COVID-19 severity correlates with airway epithelium–immune cell interactions identified by single-cell analysis,” Nat. Biotechnol., vol. 38, no. 8, pp. 970–979, Aug. 2020, doi: 10.1038/s41587-020-0602-4.

[41] C. Bleilevens et al., “Macrophage Migration Inhibitory Factor (MIF) Plasma Concentration in Critically Ill COVID-19 Patients: A Prospective Observational Study,” Diagnostics, vol. 11, no. 2, p. 332, Feb. 2021, doi: 10.3390/diagnostics11020332.

[42] J. L. Caniglia, S. Asuthkar, A. J. Tsung, M. R. Guda, and K. K. Velpula, “Immunopathology of galectin-3: an increasingly promising target in COVID-19,” F1000Research, vol. 9, p. 1078, Sep. 2020, doi: 10.12688/f1000research.25979.2.

[43] R. Kaufmann et al., “Thrombin-mediated hepatocellular carcinoma cell migration: Cooperative action via proteinase-activated receptors 1 and 4,” J. Cell. Physiol., vol. 211, no. 3, pp. 699–707, 2007, doi: 10.1002/jcp.21027.

[44] A. Gowhari Shabgah et al., “Shedding more light on the role of Midkine in hepatocellular carcinoma: New perspectives on diagnosis and therapy,” IUBMB Life, vol. 73, no. 4, pp. 659–669, 2021, doi: 10.1002/iub.2458.

[45] M. Hatakeyama, “Structure and function of Helicobacter pylori CagA, the first-identified bacterial protein involved in human cancer,” Proc. Jpn. Acad. Ser. B Phys. Biol. Sci., vol. 93, no. 4, pp. 196–219, Apr. 2017, doi: 10.2183/pjab.93.013.

