## Supplemental Figures for "Vulture: Cloud-enabled scalable mining of microbial reads in public scRNA-seq data"

### Supplementary Materials

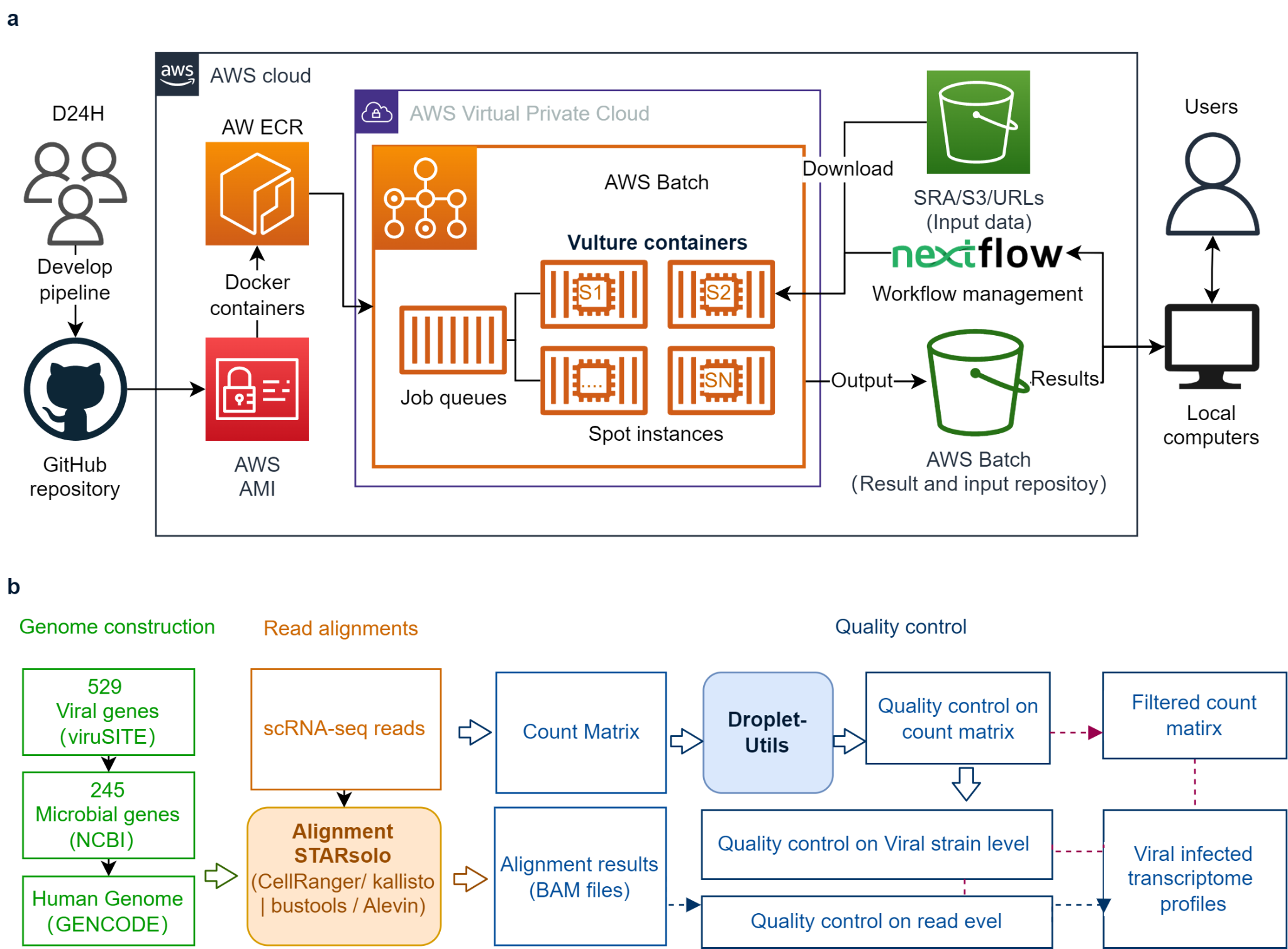

**Fig. S1:** Infrastructure of the Vulture application. **a)** Describes the cloud services application and user workflow of Vulture on the cloud. **b)** Describes the procedure of each Vulture application locally.

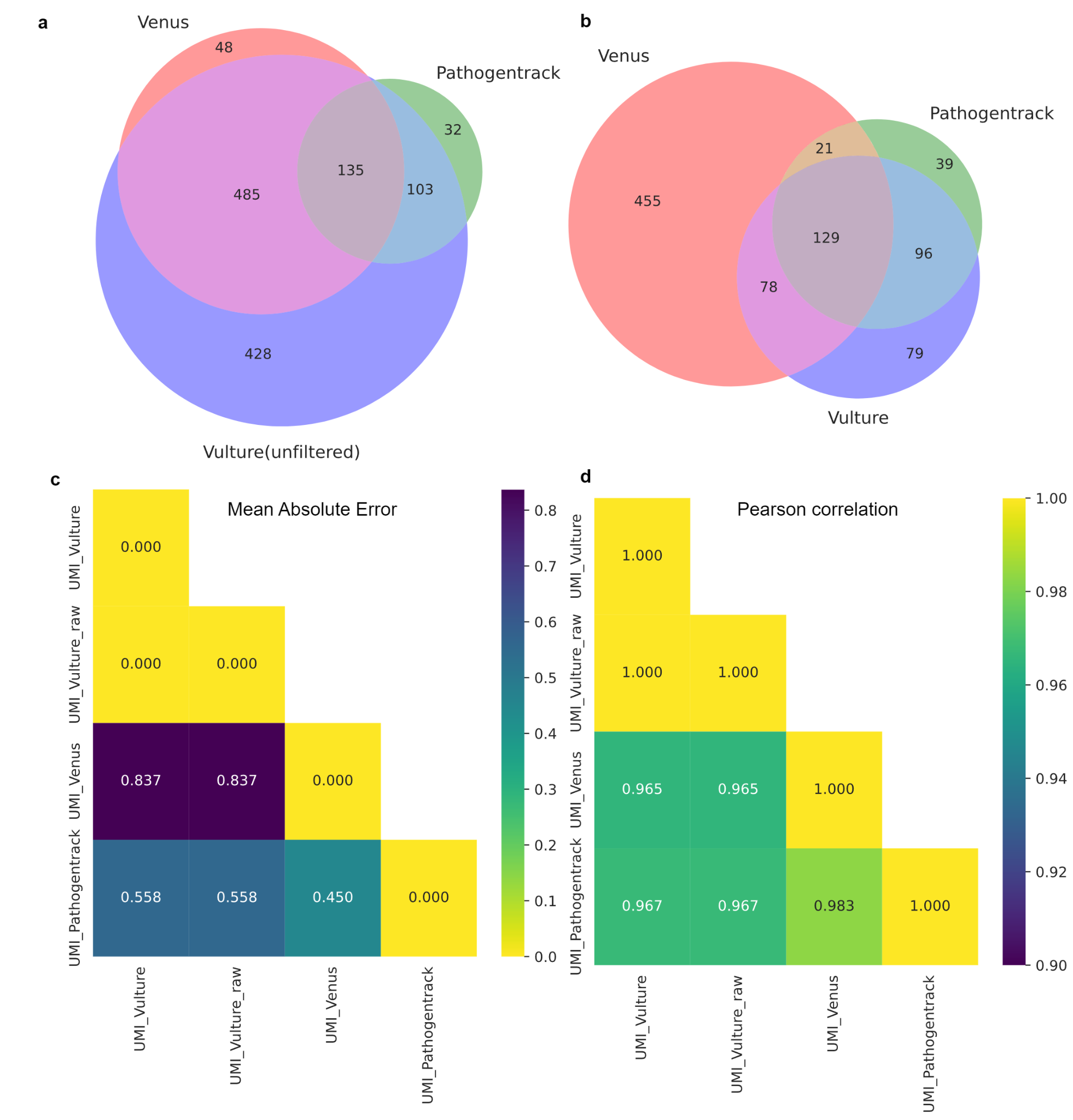

**Fig. S2** Consistency metrics among Vulture (or unfiltered) and other tested tools. **a)** and **b)** are Venn diagrams showing the interaction of SARS-CoV-2 positive cells among all tested tools. Vulture (unfiltered) is the result of Vulture before EmptyDrops filtering. **c)** is the mean absolute error (MAE) of UMIs across the intersection of SARS-CoV-2 positive cells. **d)** is the Pearson correlation of UMIs across the intersection of SARS-CoV-2 positive cells.

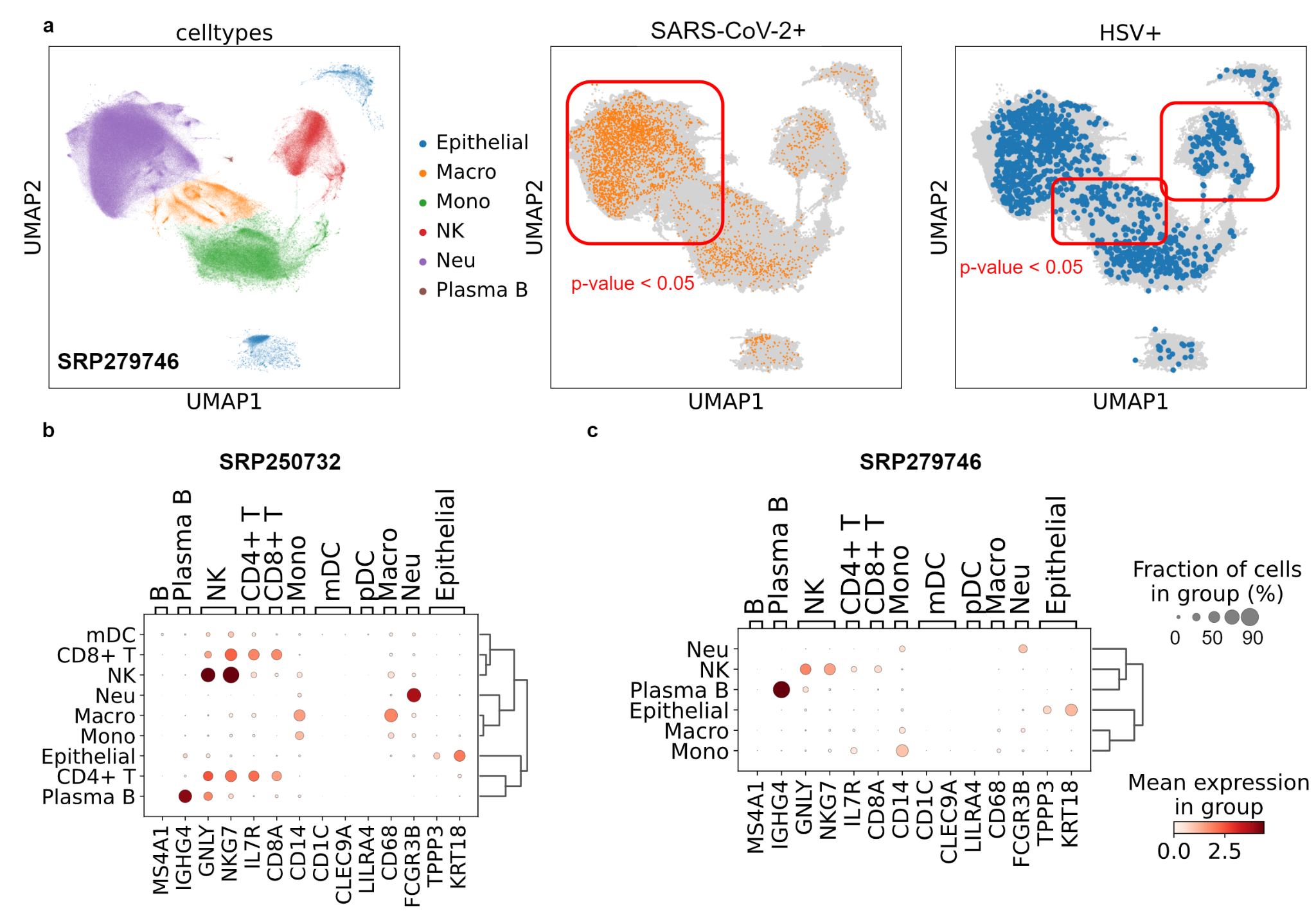

**Fig. S3** Viral calling meta-analysis results on the complete BALF sample. **a)** is a set of UMAP plots of the Bost et al. sample. The one on the left is a UMAP colored by sub-clustering annotations. The one in the middle is a plot of the SARS-CoV-2 infection. Infected cells are colored orange while other cells are gray. The one on the right is a plot of the HSV infection. Infected cells are colored blue while other cells are gray. **b)** shows the marker genes expression across cell types to identify different cell types of the Liao et al. cohort. **c)** shows the marker genes expression across cell types to identify different cell types of the Bost et al. cohort.

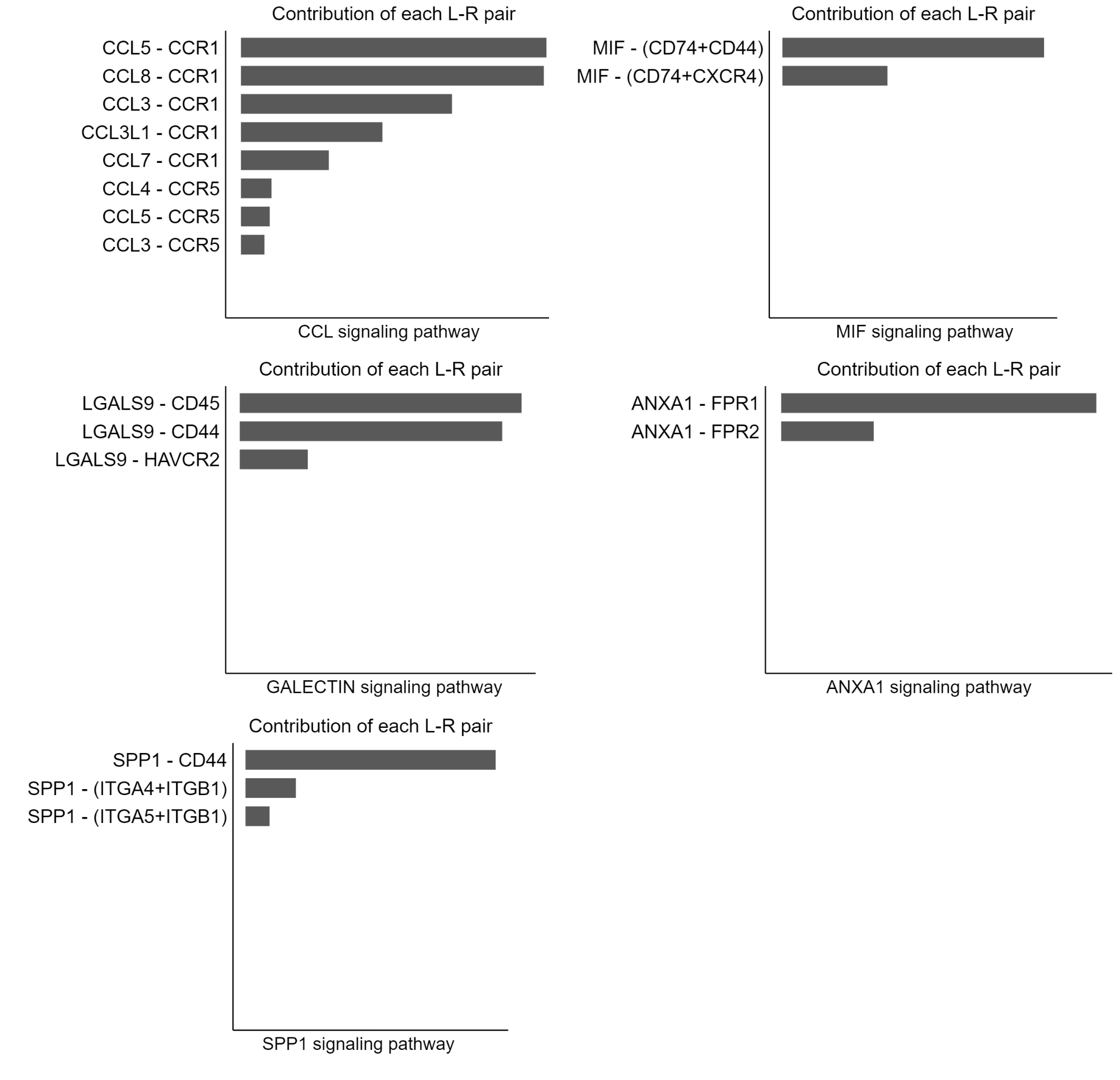

**Fig. S4** Ligand-receptor pairs with the strongest contribution among SARS-CoV-2 and hMPV-positive Macrophages and Monocytes in the COVID-19 BALF samples

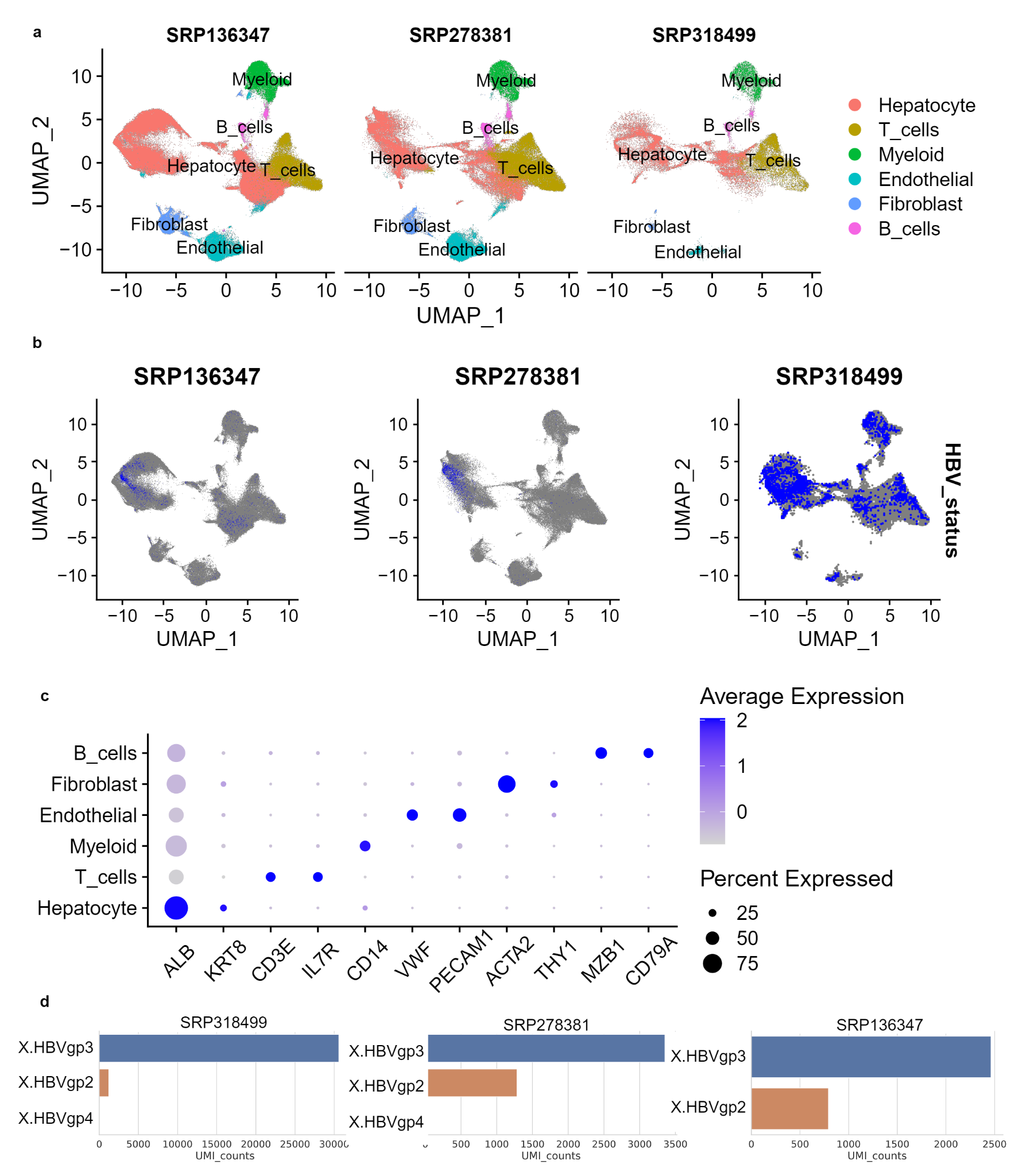

**Fig. S5** Viral calling meta-analysis results on the complete HCC sample. **a)** is a UMAP plot of the HCC sample, cells are colored by sub-clustering annotations. **b)** are UMAP plots of the HBV infection. Infected cells are colored blue while other cells are gray. **c)** shows the marker genes expression across cell types to identify different cell types. **d)** shows UMIs of the detected transcripts from HBV.

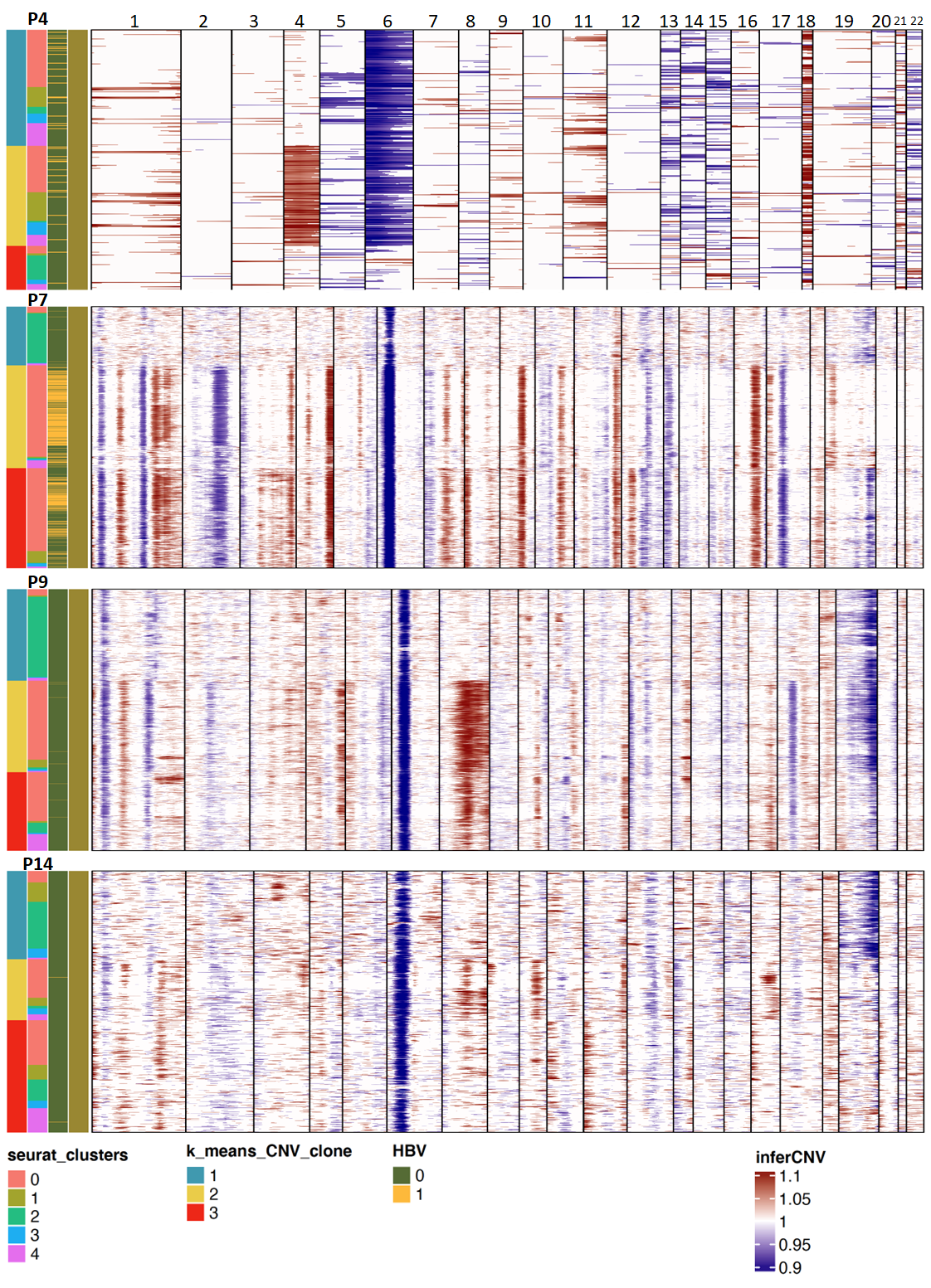

**Fig. S6** inferCNV diagram of SRP278381

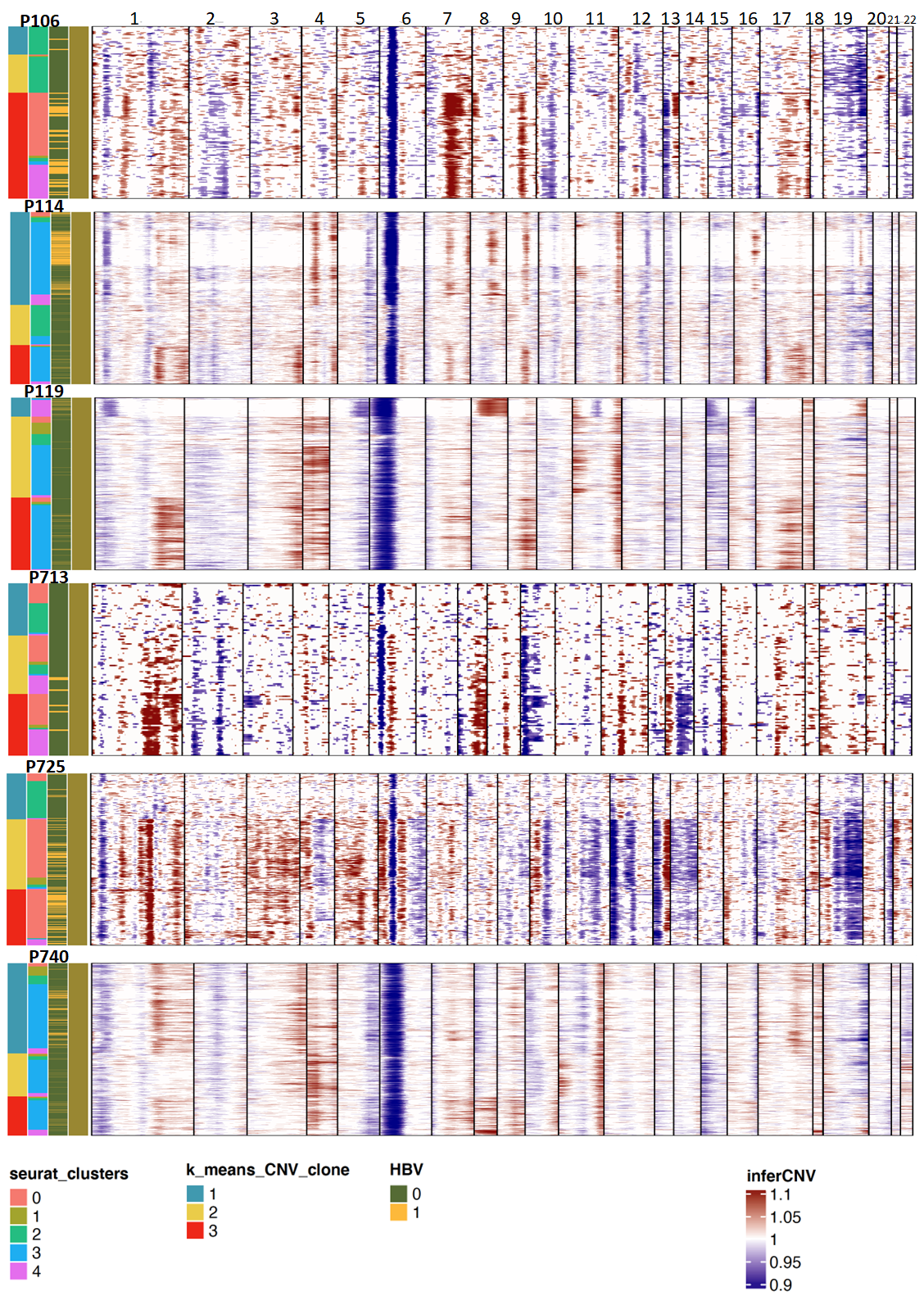

**Fig. S7** inferCNV diagram of SRP318499

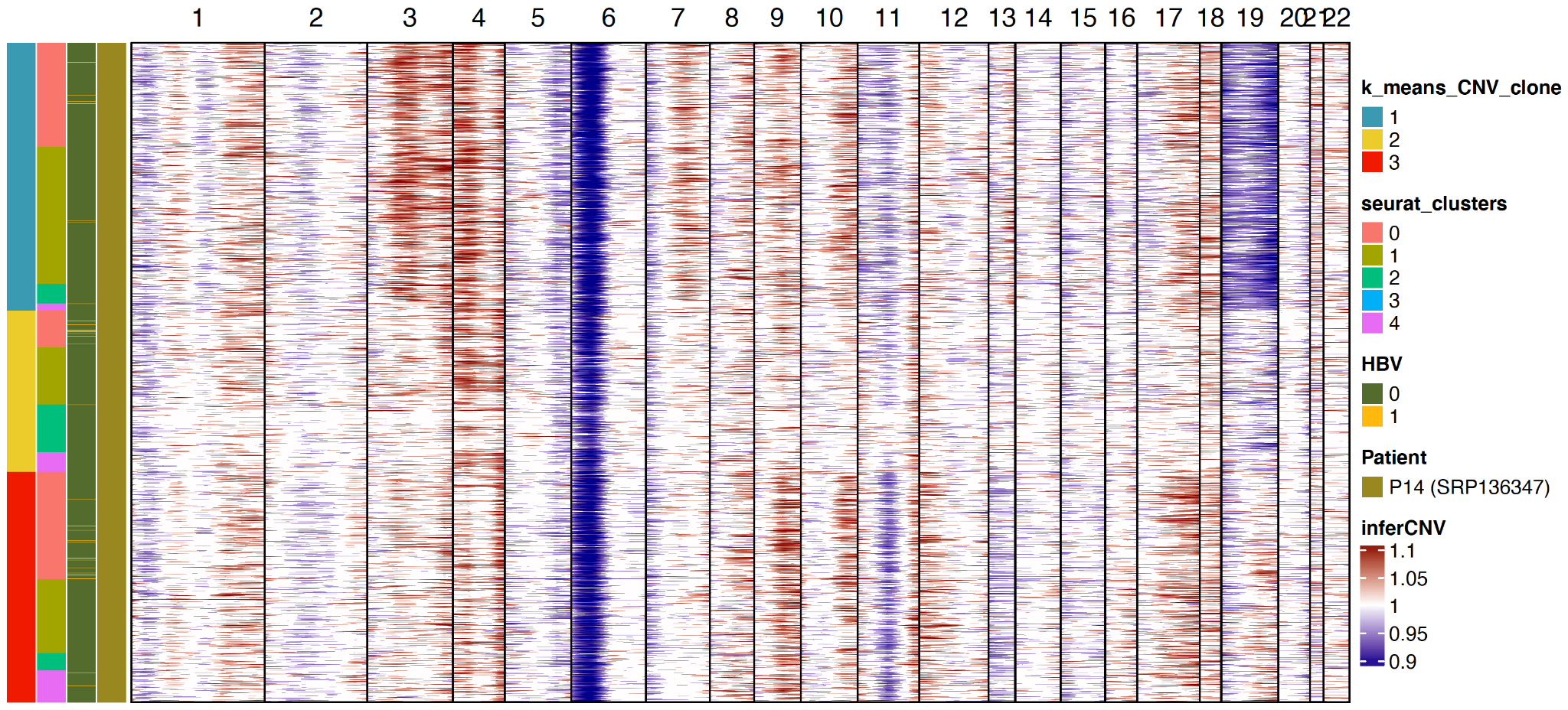

**Fig. S8** inferCNV diagram of SRP136347
